# Pacing strategy in horse racing

**DOI:** 10.1101/2020.06.11.145797

**Authors:** Quentin Mercier, Amandine Aftalion

**Affiliations:** Ecole des Hautes Études en Sciences Sociales, Centre d’Analyse et de Mathématique Sociales, CNRS UMR-8557, 54 boulevard Raspail, Paris, France

## Abstract

Thanks to velocity data on races in Chantilly (France), we set a mathematical model which provides the optimal pacing strategy for horses on a fixed distance. It relies on mechanics, energetics (both aerobic and anaerobic) and motor control. We identify the parameters useful for the model from the data. Then it allows to understand the velocity, the oxygen uptake evolution in a race, as well as the energy or the propulsive force and predict the changes in pacing according to the properties (altitude and bending) of the track.

## Introduction

Very little is known about the optimal strategy for a horse to run and win a race. In particular, due to limitations in the measurement of the mean oxygen uptake (*V̇ O*2) for a horse at high exercise, no information is available on the full *V̇ O*2 profile in a race, depending on the distance. Up to now, only measurements on treadmills have been obtained using masks [1–4]. A report on horses physiology can be found in [5]. What is known is that horses have a high aerobic capacity, about twice that of human beings, due to a high capacity for oxygen carriage and extractions, as well as a high stroke volume. They reach the maximal oxygen uptake (*V̇ O*2_*max*_) much quicker than humans, nevertheless no precise estimate of the time needed to reach a steady state *V̇ O*2 is known. Similarly, no information is available about the distance or time at which the *V̇ O*2 starts decreasing at the end of a race. Eventually, for long distances, it is not known whether the *V̇ O*2 remains almost constant for the most part of the race or oscillates around a mid value, and then at which period and at which amplitude. This paper will provide pieces of information for these issues.

Reference [6] is the only one that we know of where pacing strategy for horses is analyzed, together with the effect of drafting. Several directions of study have been investigated to better understand the effort or mechanical work developed by horses. One of them is to measure the propulsive force either on a force plate [7] or with an instrumented horse shoe [8, 9].

The aim of this paper is to provide a mathematical model able to predict the pacing strategy depending on the distance to run, the shape and topology of the track. It is based on the model developed in [10–12] for human races and it is adapted here to horses to fit the data.

Based on data provided by France Galop and the Mc Lloyd tracking device in French horse races, we are able to model the optimal horse efforts and velocity for a fixed distance, depending on the curvature and change of slopes and ramps. Our model yields in particular information on the *V̇ O*2 profile.

## Materials and methods

### Data

The data consist of two dimensional position and speed sampled at 10 Hz for horses racing in Chantilly (France), at the end of the 2019 thoroughbred horse racing season. They are provided by France Galop and are from roughly ten races. The tracking system is developed by Mc Lloyd. It is a miniaturized device which does not bring any discomfort to the horse or the jockey. Reliable data is obtained thanks to a patented positioning technology and robust mobile network data transmission, even during crowded events. The device mean accuracy is 25cm or 2 hundredth of a second. The accuracy of the system was validated on horses by comparing 1ms-accurate photo finish data on more than one hundred races with gap data on the finish line obtained after processing latitude and longitude data. The tracking system provides the latitude and longitude data sampled at 10Hz, as well as the velocity. The latitude and longitude data given by the tracker are projected over a reference track leading to the position of horses in the race.

### Model

Once we have these raw data, we have to smooth the speed using a third order Savitzky–Golay filter. Therefore, the raw data provide two curves for each horse sampled at 10Hz:

- the curve of time vs distance from start, projected on the reference track, so that each horse runs the same distance,
- the curve of velocity vs distance from start, projected on the reference track.

We use the model developed by [10–12] for the optimal strategy in running and adapt it for horses. For a fixed distance, this model predicts the final time, the velocity curve and the effort developed by the horse to produce the optimal strategy. It depends on the geometry of the track, the ramps and slopes and the physiology of the horse. This physiology is taken into account through a number of parameters that have to be identified numerically for each horse thanks to the data:

- the maximal propulsive force per unit of mass *f_M_*,
- the global friction coefficient *τ* which encompasses all kinds of friction, both from joint and track. In total, *f_M_ τ* is the maximal velocity,
- the maximal decrease rate and increase rate of the propulsive force which is related to the motor control of the horse: a horse, like a human being, cannot stop or start its effort instantaneously, but needs some time or distance to do it. This is what our control parameters *u*_−_ and *u*_+_ will provide,
- the total anaerobic energy or maximal accumulated oxygen deficit *e*^0^,
- the *V̇ O*2 profile as a function of distance, namely the distance *d*_1_ at which the maximum of *V̇ O*2 is reached, the distance *d*_2_ at which *V̇ O*2 decreases, and the relative decrease with respect to the maximum value. This is a curve *σ*(*s*) where *s* is the relative distance from start, but in fact, in the model, it is a curve *σ*(*e*(*s*)) where *e*(*s*) is the remaining anaerobic energy. The profile of *σ* is to be identified from the data.

For fixed values of these parameters, the model of [10–12] adapted to horses yields an optimal control problem based on a system of coupled ordinary differential equations for the instantaneous velocity *v*(*s*), the propulsive force per unit of mass *f* (*s*), the anaerobic energy *e*(*s*), where *s* is the distance from start. The system relies on Newton’s second law of motion taking into account the positive slope or negative ramp *α*(*s*), the energy balance (between the aerobic contribution *V̇ O*2 or *σ*(*e*), the anaerobic contribution *e*(*s*) and the power developed by the propulsive force) and finally the control on the variations of the propulsive force. A crucial piece of information to be taken care of is the centrifugal force in the bends. This force does not act as such in the equation of movement but limits the propulsive force through a constraint which yields a decrease in the effective propulsive force in the bends.

For the ease of completeness, we provide below the full optimal control problem though the results of the paper do not require to understand it and the reader can skip this paragraph as a first reading. Instead of writing the equations of motion in the time variable, we write them using the distance from start *s*. This amounts to dividing by *v* the derivatives in time in order to get the derivatives in space. We also write the equations per unit of mass. Let *d* be the length of the race and *g* = 9.81 the gravity. Let *c*(*s*) denote the curvature at distance *s* from the start, which is provided for each track and *α*(*s*) be the slope coefficient. Let *v*^0^ be the initial velocity. Let *e*^0^, *f_M_*, *τ*, *u*_−_, *u*_+_ and the function *σ* be given. They are identified for a specific race and horse. The optimal control problem (where minimizing the final time is equivalent to minimizing the integral of the inverse of the velocity) coming from the Newton law of motion and the energy conservation is

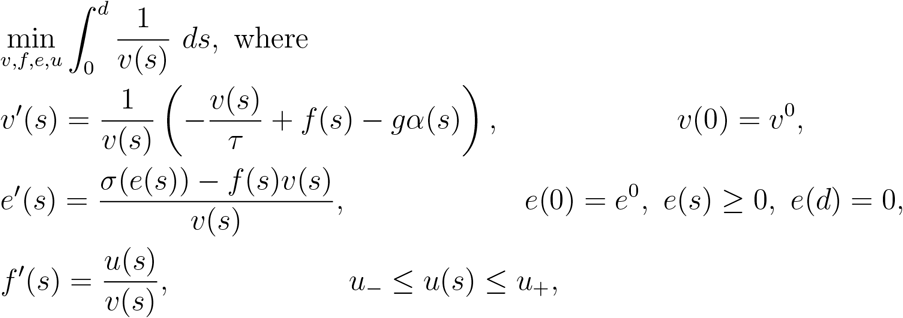

and under the state constraint

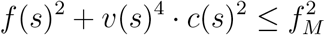

for *s* ∈ [0, *d*].

The optimal control problem for horse performance is solved using Bocop, an open licence software developed by Inria-Saclay France [13]. This yields the optimal strategy depending on the length of the race, including the velocity profile and the *V̇ O*2 profile. Our aim is therefore to identify the physiological parameters *f_M_*, *τ*, *u*_+_, *u*_−_, *e*^0^, *σ*(*e*) from the available data.

### Identification process

The identification process is made through a bi-level optimization procedure looking for minimizing errors between the response of a Bocop simulation and the data through the following objective

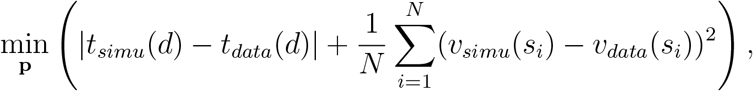

where the subscript *simu* (resp. *data*) refers to the variable extracted from the simulation (resp. data). The distance *s*_1_ = 0 is the beginning of the race and *s_N_* = *d* is the length of the race, while *s_i_* are intermediate distances. The parameter **p** refers to the vector containing all the variables to identify:

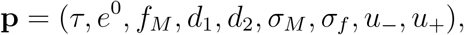

where *σ_M_* is the maximal value of *σ* and *σ_f_* its final value. The objective is made up of two parts: first the difference in final time at the end of the race *d*, and then the mean square error over the speed measured at *N* points. For our identification process, *N* is taken equal to one thousand and the points are evenly distributed between *s*_1_ and *s_N_*.

The algorithm used here is a particle swarm optimization method [14] available in the pyswarm library in Python 3, part of the family of the heuristic optimization methods [15]. The main advantage of such a method is its good ability to explore the design space and its ease of use and implementation. A swarm of designs {**p**_*i*_} is tested, that is the optimal control problem is solved for these parameters using Bocop, and its objective value is calculated. The performance influences the speed and the direction of the particles inside the design space for the next iteration. At the end of the process, the best score particle is kept as the result. The stopping criterion of the algorithm is set such that the algorithm is unable to find new particles for which the objective is at least 10^−7^ better than the best score observed until then. For all the examples treated in this paper, the swarm size is set to 50 particles and a maximum of 150 iterations. Each identification process has reached the stopping criterion described. This method is well suited to our problem since the objective space has a lot of local minimizers so that gradient based methods can get stuck in local minimizers. It insures overall robustness in the results as the design space is always well explored before converging toward a particular area of the design space.

### Topology of the tracks

We have studied three types of races: a 1300 meters, a 1900 meters and a 2100 meters in Chantilly. The GPS track is shown in Fig 1. The 1300m starts with a straight, then there is a bend before the final straight. The 1900m starts earlier with an almost straight and follows the 1300m. The 2100m starts in the final straight of the other races, makes a closed loop before reaching the same final straight and finish line.

**Fig 1.**
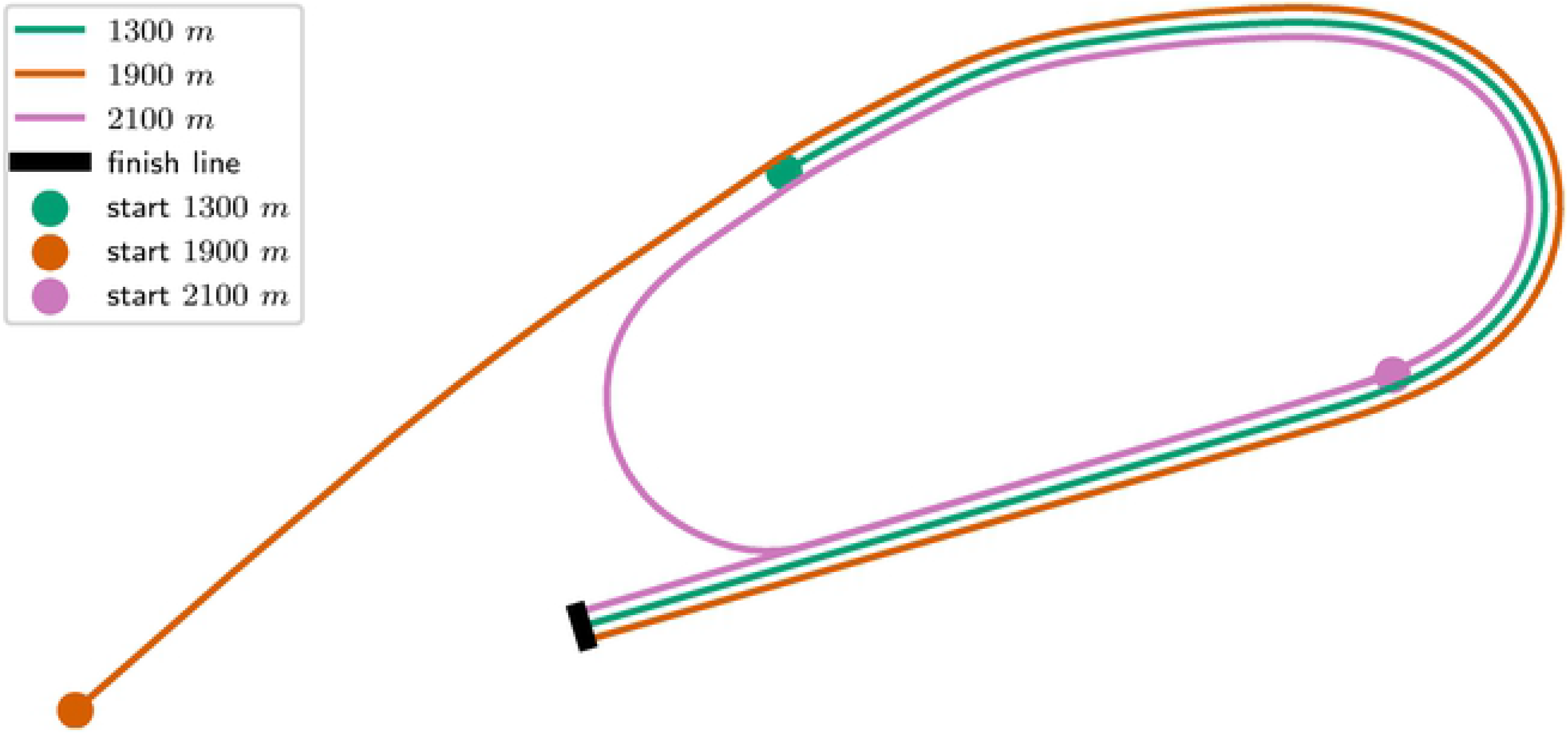
GPS track of the 1300m, 1900m and 2100m in Chantilly, France.

The elevation and curvature profiles are provided by France Galop and illustrated in Fig 2. The tracks are made up of straights (zero curvature), arcs of circles (constant curvature) and clothoids (curvature increasing linearly with distance). A clothoid is the usual way to match smoothly a straight and an arc of a circle since the curvature increases linearly. It is used for train tracks and roads as well. It allows smooth variations of velocities which are more comfortable for horses.

**Fig 2.**
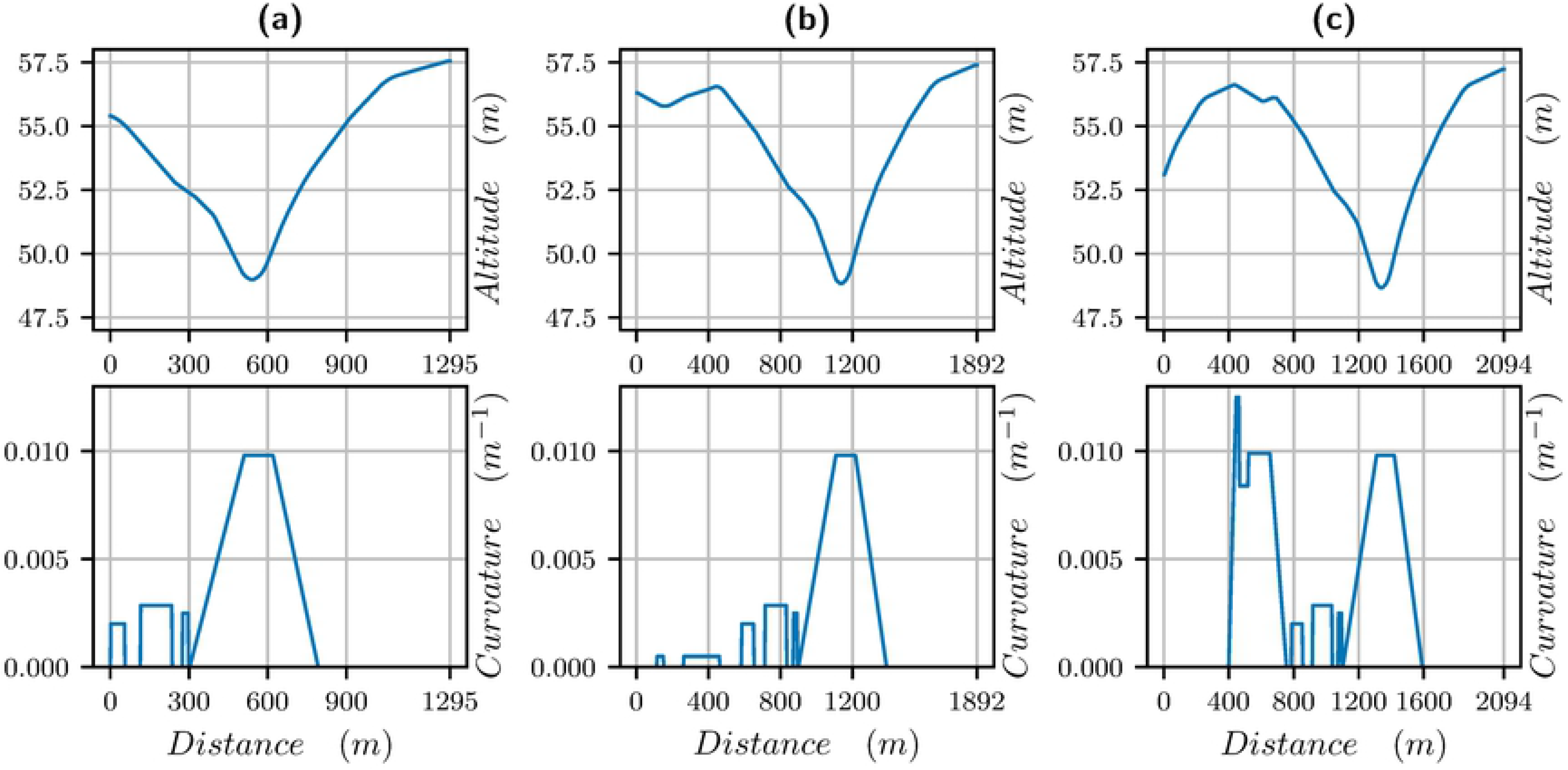
Altitude in meters (first line) and curvature (second line) vs distance for different tracks: (a) 1300m (first column), (b) 1900m (second column) and (c) 2100m (third column). The last 1300m are always the same.

The specificity of the track is that there is a bend of 500 meters before the final straight, where the track is going down in the first quarter (about 1.5%) and going up in the second quarter (about 2%). In the 2100 meters, there is also a bend of 400 meters just after the start, which is first going up and then down. We will see that curvature and altitude have a strong effect on the pacing.

Let us point out that the track is banked but, because data correspond to horses close to the inner part of the track, the banking is not meaningful for the data and will not be taken into account here.

## Results

We have chosen three significant races of 1300m, 1900m and 2100m. For each one, we have taken the data of a horse which seems to have run an optimal race. We describe below the results of our simulations.

### 1300 meters

The parameters identified for this race are in Table 1. The velocity data (raw and smoothed) and the velocity computed with our model are plotted in Fig 3 for the 1300m. We observe a very good match between the curves: there is a strong start with the maximal velocity being reached in 200 meters. Then the velocity decreases, and in particular in the bend, between 300 and 600 meters from start. Though the track is going down, the centrifugal force reduces the propulsive force as we see in Fig 4b (the black curve shows the limitation due to the centrifugal force). It is only when reaching the clothoid, before the final straight, after 600 meters, that the horse can speed up again. The end of the race is uphill and the velocity decreases though the horse reaches the straight. Nevertheless, a decrease in velocity at the end of such a race takes place even on a flat track.

**Table 1.** Identified parameters for the 1300m

| $\tau$ | $e^0$ | $f_M$ | $d_1$ | $d_2$ | $\sigma_M$ | $\sigma_f$ | $u_-$ | $u_+$ |
| --- | --- | --- | --- | --- | --- | --- | --- | --- |
| 3.911 | 2731 | 5.150 | 421.5 | 559.9 | 47.0 | 40.71 | -1.693e-03 | 1.504e-03 |

**Fig 3.**
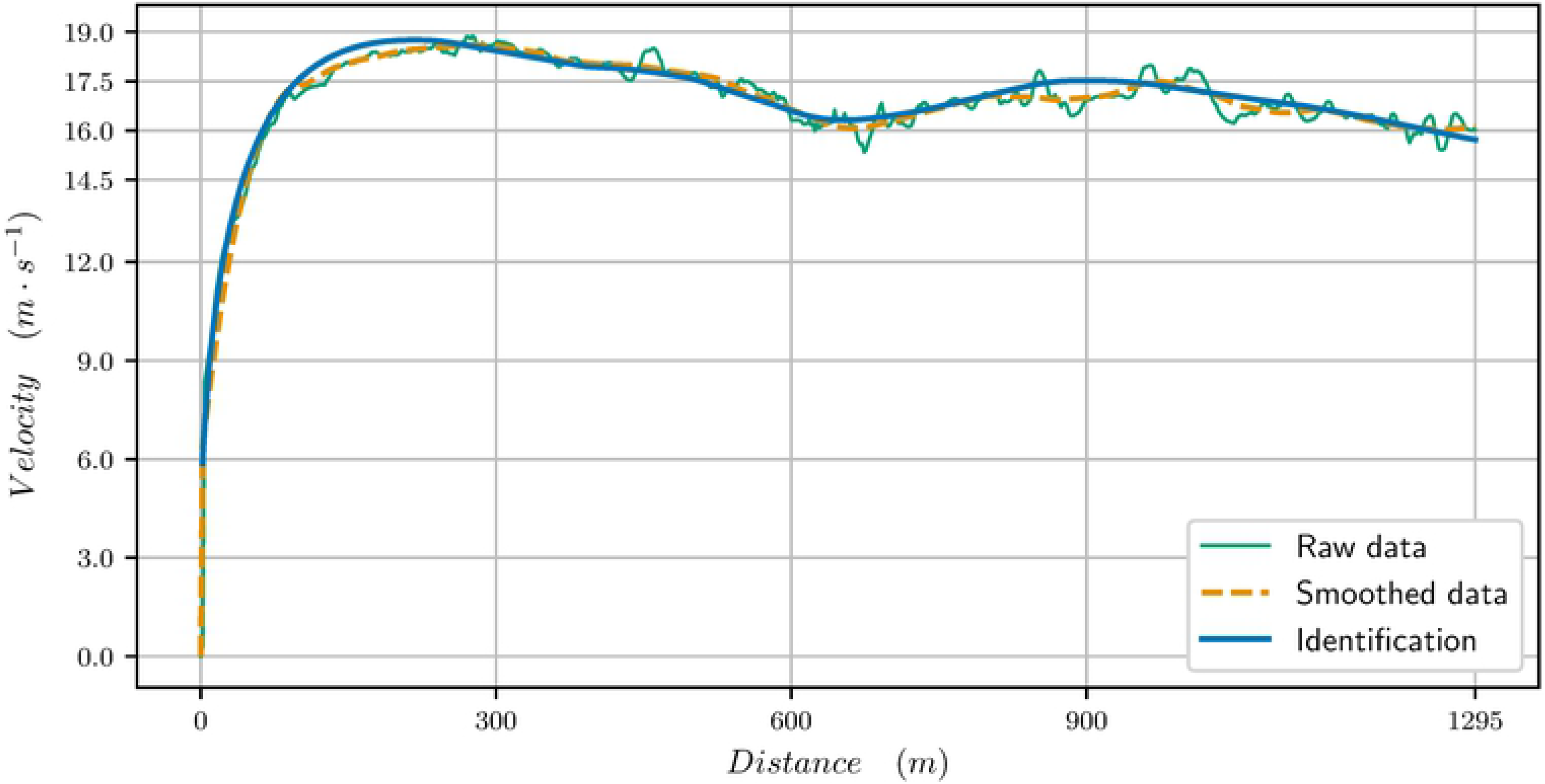
Velocity data (raw data and smoothed data, *t_f_* = 76.544s) and computed velocity for the identified paramaters (*t_f_* = 76.508s) for the 1300 meters race

**Fig 4.**
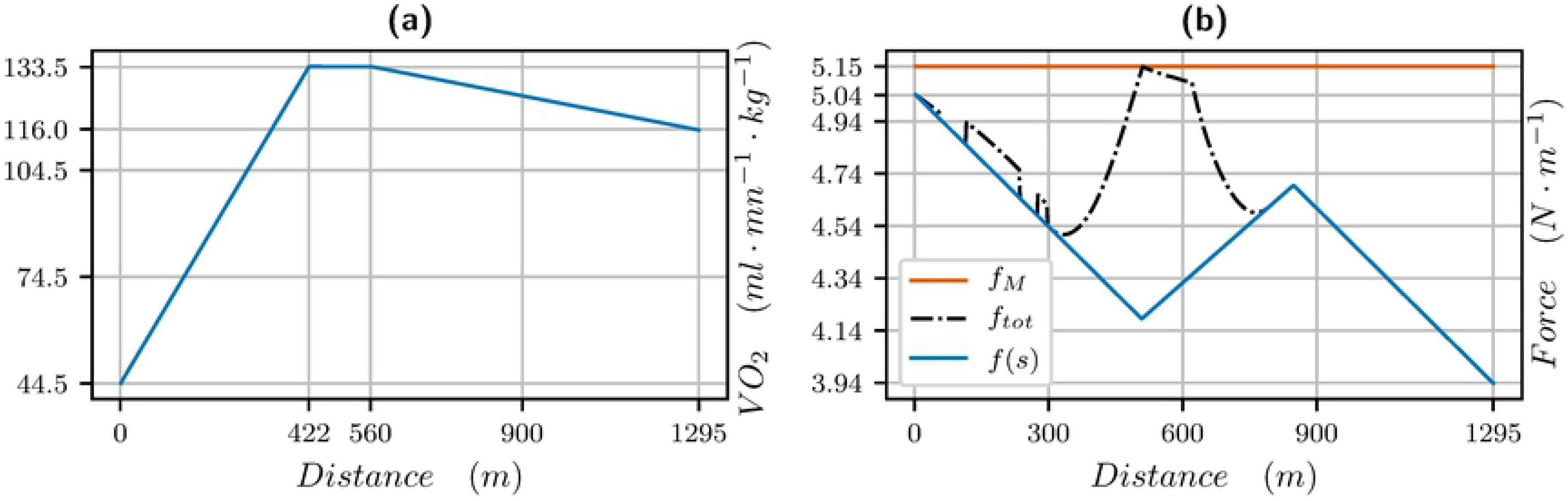
*V̇ O*2 (left) and propulsive force (right) vs distance for a 1300 meters: blue is the propulsive force *f* (*s*) in the direction of movement, black is the effective propulsive force 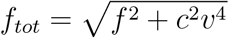 taking into account the centrifugal force, where *c* is the curvature.

The *V̇ O*2 curve vs distance and propulsive force vs distance are plotted in Fig 4. We measure *σ* in *J/s/kg* but we want to plot the results in terms of *ml/mn/kg* knowing that one liter of oxygen produces roughly 21 *kJ* (see [16] and also [17]), so we have to multiply our data for *σ* by 60*/*21. For a maximal value of *σ* equal to 47, this yields a *V̇ O*2_*max*_ of 133.6 *ml/mn/kg*. We see that the *V̇ O*2 is increasing for about 400 meters, while the force is decreasing. Then when the *V̇ O*2 decreases, the force and thus the velocity increase until 900 meters when the slope and end of race lead to a decrease of force and velocity. We point out that the value of the propulsive force is higher than the ones found in [8, 9] but the velocity is also much higher.

### 1900 meters

The parameters identified for this race are in Table 2. The velocity data (raw and smoothed) and the velocity computed with the model are plotted in Fig 5 for the 1900 meters. We observe that there is a strong start with the maximal velocity being reached in 300 meters. Then the velocity decreases. Between 900 and 1400m, we see the effect of the bend: at the beginning of the bend, the track is going down and the horse slightly speeds up; then the centrifugal force reduces the velocity but the velocity increases again at the end of the bend. The end of the race is with a strong acceleration before the final slight slow down. The *V̇ O*2 curve vs distance and propulsive force vs distance are plotted in Fig 6. We see that *V̇ O*2 is increasing for about 400 meters, while the force starts at maximal value. Then, the *V̇ O*2 is constant, the force and the velocity decrease to a mean value. At the end, the *V̇ O*2 decreases when the residual anaerobic energy reaches a third of its initial value. The effect of the bend and centrifugal force are obvious: it leads to a decrease in propulsive force and velocity. We see on the force profile that there is a very strong acceleration in the end. It can only take place after the bend where the centrifugal force reduces the available propulsive force.

**Table 2.** Identification parameters for the 1900 meters

| $\tau$ | $e^0$ | $f_M$ | $d_1$ | $\sigma_M$ | $d_2$ | $\sigma_f$ | $u_-$ | $u_+$ |
| --- | --- | --- | --- | --- | --- | --- | --- | --- |
| 3.73 | 2702 | 4.74 | 379 | 54.6 | 1392 | 40.3 | -3.69e-03 | 3.94e-03 |

**Fig 5.**
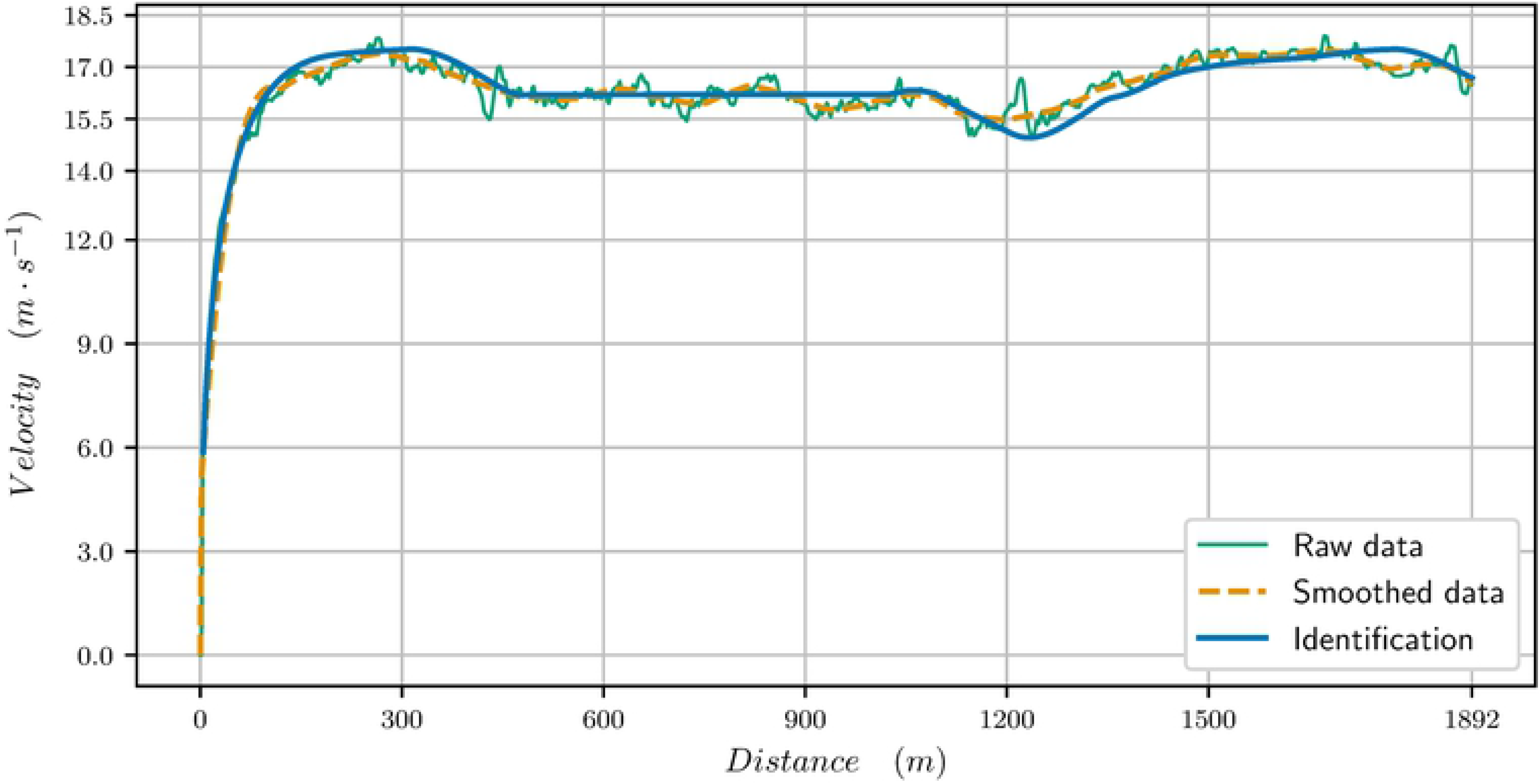
Velocity data (raw and smoothed, *t_f_* = 116.460s) and computed velocity (*t_f_* = 116.460s) for the identified parameters for the 1900 meters race

**Fig 6.**
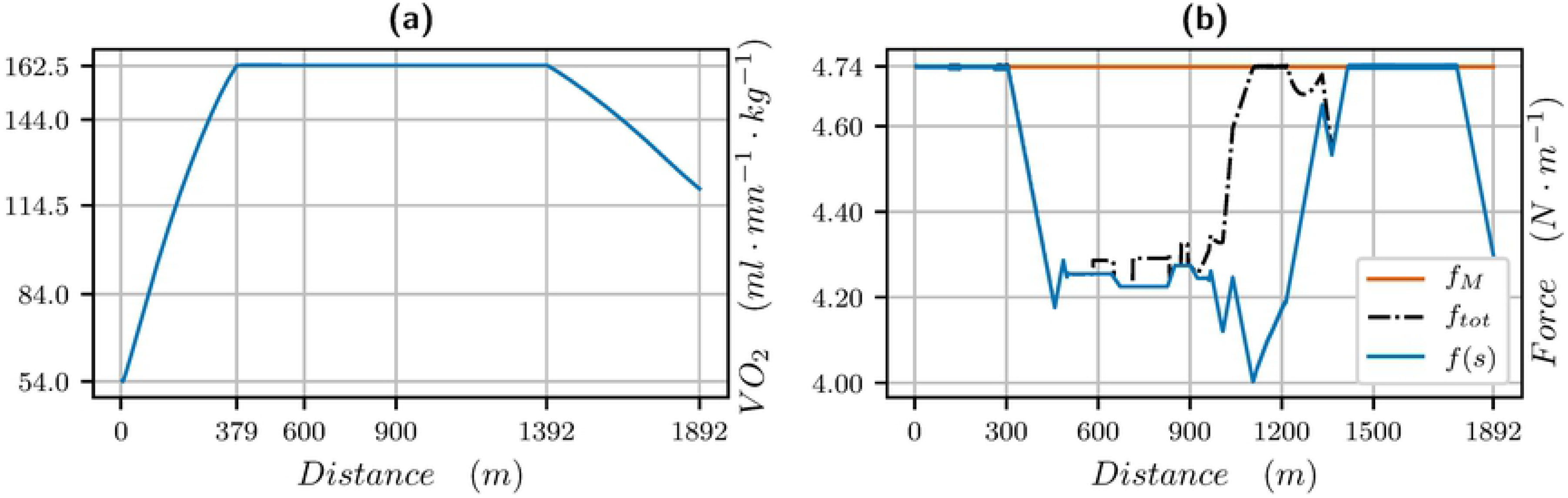
*V̇ O*2 (left) and propulsive force (right) vs distance for a 1900 meters: blue is the propulsive force *f* (*s*) in the direction of movement, black is the effective propulsive force 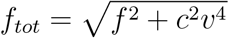 taking into account the centrifugal force, where *c* is the curvature.

In Fig 7, we have plotted a zoom on the velocity curve for the identified parameters, and then have removed the effect of the slope (flat track), of the curvature (straight track) and of both (flat, straight track). This allows to notice the specific effects: a bend reduces strongly the velocity (pink curve); on the real track (brown curve), because the first part of the bend is going down, the reduction in propulsive force and velocity is not so strong. The end of the track is uphill and one notices that the velocity curves corresponding to going up (brown, red) cannot provide a speeding up as high as the two others. The combination of slopes and ramps of this track (red curve) reduce the velocity and final time in total though the profile is very similar. We also point out that though these are local effects, they have a global influence on the strategy since they change the mean velocity.

**Fig 7.**
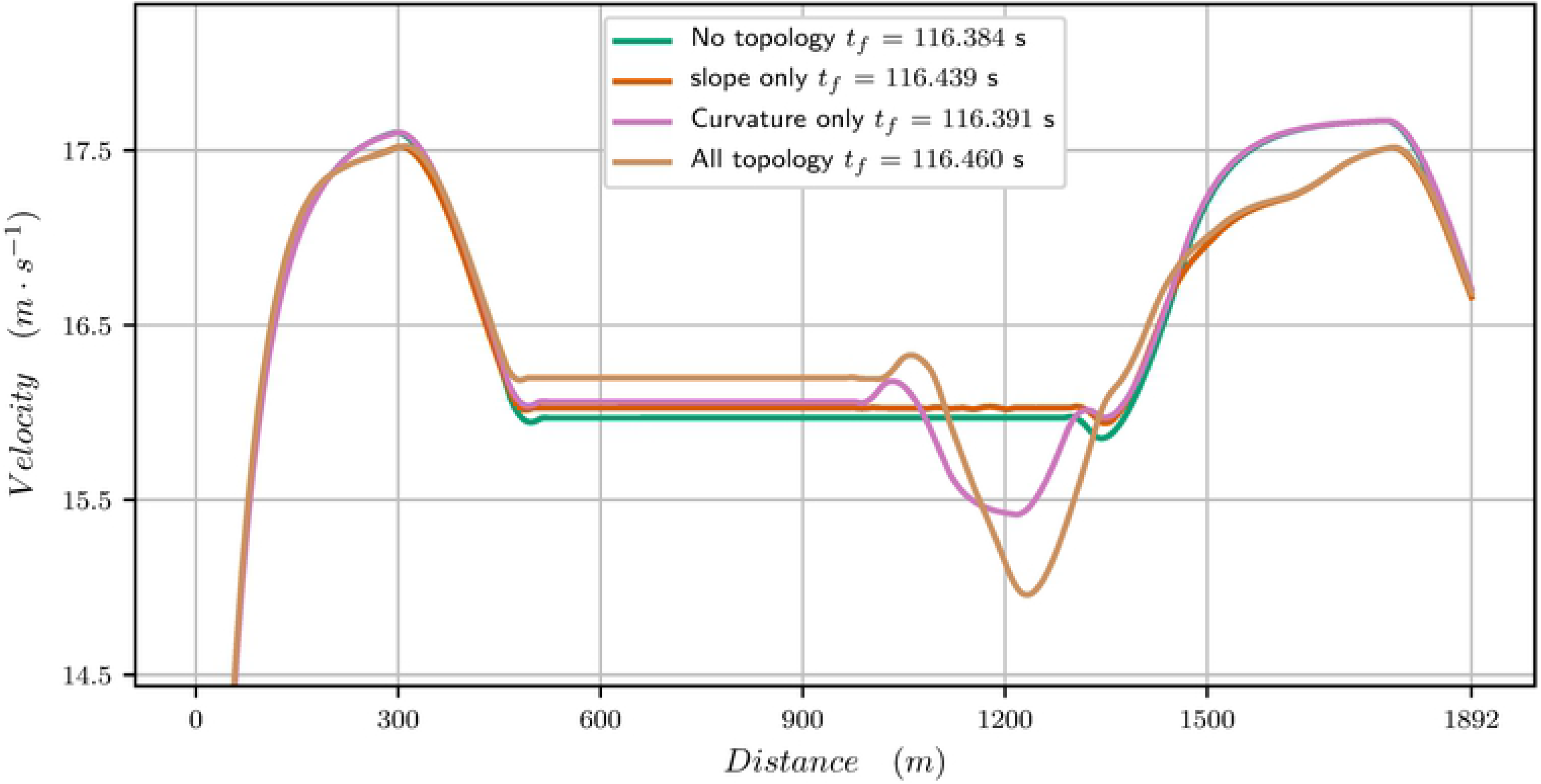
Effect of the slope and curvature on the velocity curve (zoom) for the 1900 meters. Brown is the velocity curve of the race, red with the slope only (straight track), purple with curvature only (flat track) and green is a flat, straight track.

### 2100 meters

The parameters identified for this race are in Table 3. The velocity data and the computed velocity are plotted in Fig 8 for the 2100 meters. The *V̇ O*2 and propulsive force are plotted in Fig 9. Here, the *V̇ O*2 is modified to match the behaviour observed in [18] for humans where the *V̇ O*2 first reaches a peak value, before the mean race value. The first bend going up requires a rise of *V̇ O*2 at the beginning of the race. We observe that there is a strong start with the maximal velocity being reached in 200 meters. Then the velocity decreases and reaches a plateau. This plateau has been analyzed for human race in [19] and is related to a turnpike phenomenon. It is very likely that the horse pacing strategy for long races can be analyzed with this mathematical tool as well.

**Table 3.** Identification parameters for the 2100 meters

| $\tau$ | $e^0$ | $f_M$ | $d_1$ | $\sigma_1$ | $d_2$ | $\sigma_M$ | $d_3$ | $\sigma_f$ | $u_-$ | $u_+$ |
| --- | --- | --- | --- | --- | --- | --- | --- | --- | --- | --- |
| 3.637 | 3567 | 5.50 | 304 | 51.29 | 524 | 46.71 | 1613 | 41.67 | -1.928e-03 | 9.922e-04 |

**Fig 8.**
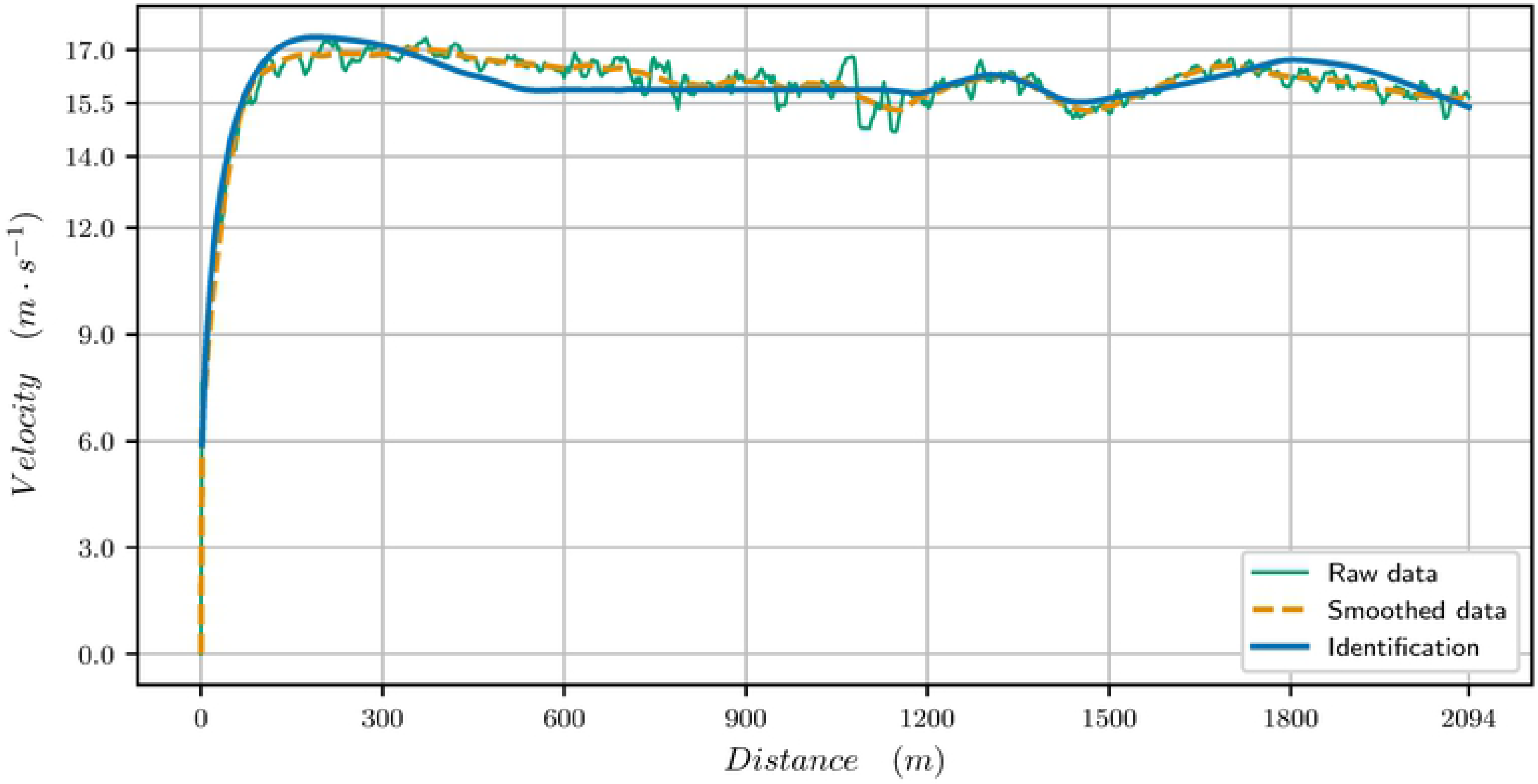
Velocity data (raw and smoothed *t_f_* = 130.933s) and computed velocity (*t_f_* = 130.97s) for the identified parameters for the 2100 meters race

**Fig 9.**
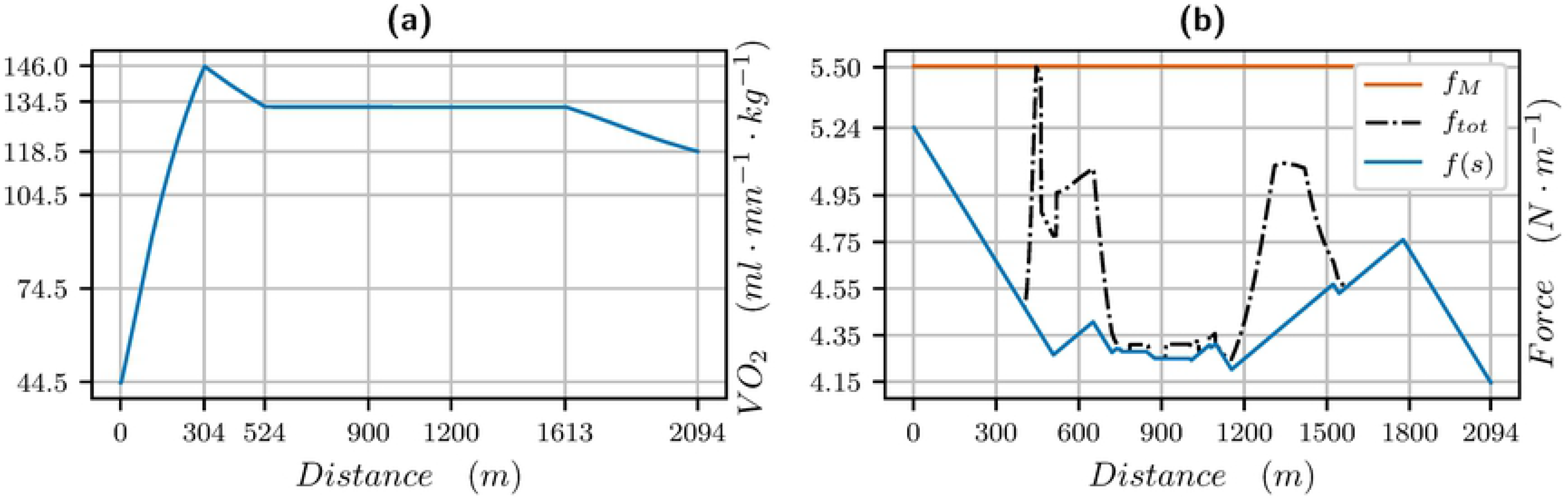
*V̇ O*2 (left) and propulsive force (right) vs distance for a 2100 meters: blue is the propulsive force *f* (*s*) in the direction of movement, black is the effective propulsive force 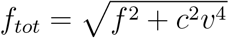 taking into account the centrifugal force, where *c* is the curvature.

The first bend has a strong curvature and therefore reduces drastically the velocity as we can see in Fig 9b: the propulsive force is reduced in the first bend. In the last bend, as in the previous race, the velocity decreases and increases again at the end of the bend. The end of the race is similar to the 1300 meters, with a strong acceleration before the final slow down. The horse in this race is not as good in terms of performance as the one in the 1900m and he cannot maintain his velocity similarly at the end of the race.

## Discussion

### Results on *V̇O*2

From experiments on human races [18, 20], it is expected that the *V̇ O*2 curve vs distance is

- increasing to a maximal value and then decreasing for short exercises,
- increasing and reaching the maximal value *V̇ O*2_*max*_, and then decreasing at the end of the race when the residual anaerobic energy is less than 30%,
- reaching a peak value which is higher than the value along the race for moderate length exercises.

Our simulations and identifications yields that the behaviour is the same for horses. The results of our simulations even provide precise information on the *V̇ O*2 curve all along the race. We see in Fig 4a, 6a, and 9a, that

- the maximal value of *V̇ O*2 is reached in about 400m, that is about 20 to 30 seconds from start, which is indeed much quicker than humans, (this is consistent with [1]),
- the 1300 meters is a short exercise where *V̇ O*2 increases and decreases,
- for the 1900 and 2100 meters race, the *V̇ O*2 reaches a mean value during the race and decreases about 500 meters from the finish line, when, as for humans, the residual anaerobic energy is about a third of the initial value. The launch of the sprint is optimal when the *V̇ O*2 starts decreasing at the end of the race, and this follows from optimal control theory. Numerically, we can observe that the decrease in *V̇ O*2 and increase in velocity at the end of the race take place at the same time,
- in a 2100 meters race, the *V̇ O*2 reaches a peak value before the mean value of the race.

The longer the horse can maintain its maximal value *V̇ O*2_*max*_, the better the performance is. Because the change of slope in *V̇ O*2 is related not exactly to the distance but to the available stock in anaerobic capacity, a high anaerobic capacity is all the more important to maintain high velocities all along the race. A strong acceleration at the beginning of the race allows to reach the maximal value of *V̇ O*2 quickly and is the best strategy. Jockeys often start slower than the optimal strategy being afraid that if the horse accelerates too strongly with respect to its capacity at the beginning of the race, then the drop in velocity at the end of the race will be bigger.

It is very likely that for horses, as for humans and explained in [18], results on treadmills are very different from measurements during a race. Indeed, as pointed out in [18], exercise at constant velocity yield contradictory results with exercises in a real race in terms of *V̇ O*2 or pacing strategy. On a treadmill, the *V̇ O*2 increases to reach a maximum value, whereas on a real race a decrease of *V̇ O*2 is observed. In many cases, the best performance is achieved with a fast start, when the pace at the beginning of the race is higher than the pace at the end. While a fast start helps to speed up *V̇ O*2 kinetics, and limits the participation of the anaerobic system in the intermediate part of the race, nevertheless, if it is too fast, it has the potential to cause fatigue and have an overall negative effect on the performance. Therefore, a departure which is too fast with respect to the horse’s capacity increases the participation of the anaerobic system at the beginning of the race and can damage the final performance. Eventually, a fast start does not necessarily induce a good performance. But a high value of *vV̇ O*2_*max*_ can allow a faster start velocity without increasing the *O*_2_ deficit.

As soon as there is a slope or ramp, the *V̇ O*2 is also impacted. Here, in Chantilly, the slope coefficients are strong enough to change the pacing strategy but not strong enough to modify the overall *V̇ O*2 profile. The only effect is on the pacing strategy. For stronger slopes, one would need to take into account additionally that *V̇ O*2 increases with slopes both downhill and uphill as explained in [6] and [21].

### Energy

Horses have two distinct types of energy supply, aerobic and anaerobic. In our model, it is *e*^0^ which estimates the anaerobic energy supply. For human races, recent research suggest that energy is derived from each of the energy-producing pathways during almost all exercise activities [22], which is what we also observe in our simulations.

In Table 4, we have computed the percentage of anaerobic energy to the total energy according to the length and duration of the race. The horse of the 1900m race has a very strong *V̇ O*2_*max*_ and therefore uses a lower anaerobic energy. We point out that the values estimated in [1] on a treadmill seem to be under estimated with respect to ours: for an exercise of duration 130 seconds, they find an anaerobic contribution of around 30%, which is smaller than our value. Indeed, in a race, for a similar duration of exercise, velocities are much higher than on a treadmill, leading to a bigger contribution of the anaerobic supply [22].

**Table 4.** Percentage of anaerobic contribution in the total energy during the race

| Race length (m) | duration (s) | $\dot{V}O_2$ deficit |
| --- | --- | --- |
| 1300 | 76 | 47.32% |
| 1900 | 116 | 32.09% |
| 2100 | 131 | 38.01% |

### Effect of slopes, ramps and bends on a race

As evident from the data of [23], on a tight bend, horses slow down a lot. In Chantilly, the bends have a radius of at most 100m, which is not tight, but still has a strong impact on the pacing strategy.

To better illustrate this effect, we choose a race of 1900 meters and set an imaginary slope or ramp or bend for one third of the race at the beginning, middle or end of the race (that is roughly 630m) with the following configuration:

- either a positive slope of +3% for 630m,
- or a negative slope of −3% for 630m,
- or an arc of circle with a curvature of 1*/*100*m^−^*^1^ for 630m.

Fig 10, 11 and 12 provide the optimal velocity vs distance for the 1900 meters parameters. The common feature is that a local change of elevation or bend does not only produce a local change in velocity but changes drastically the whole velocity profile and mean value.

**Fig 10.**
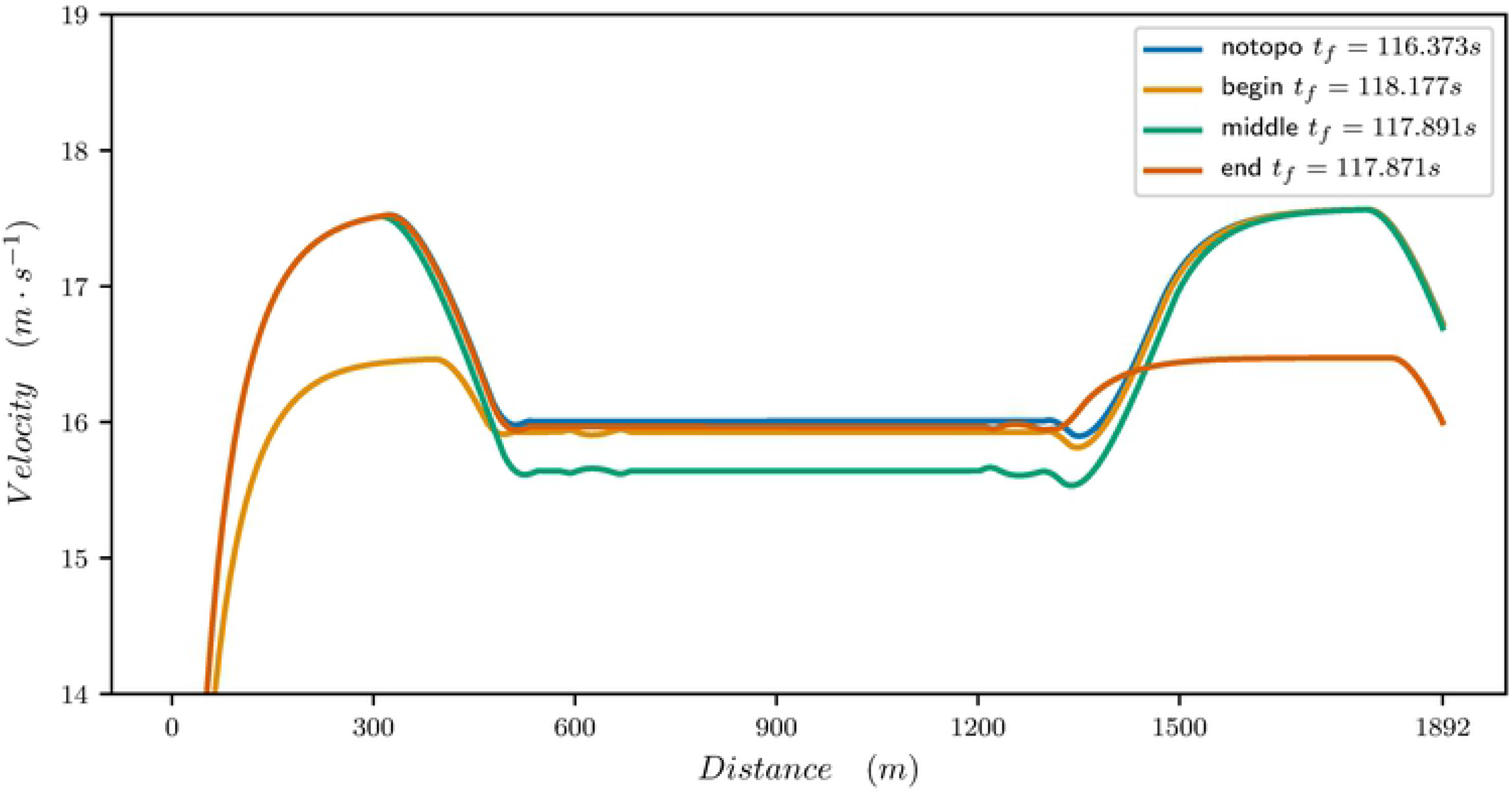
Optimal velocity profile for a 1900m: flat straight track (blue), +3% slope for 630m at the beginning of the race (orange), +3% slope for 630m at the middle of the race (green), +3% slope for 630m at the end of the race (red).

**Fig 11.**
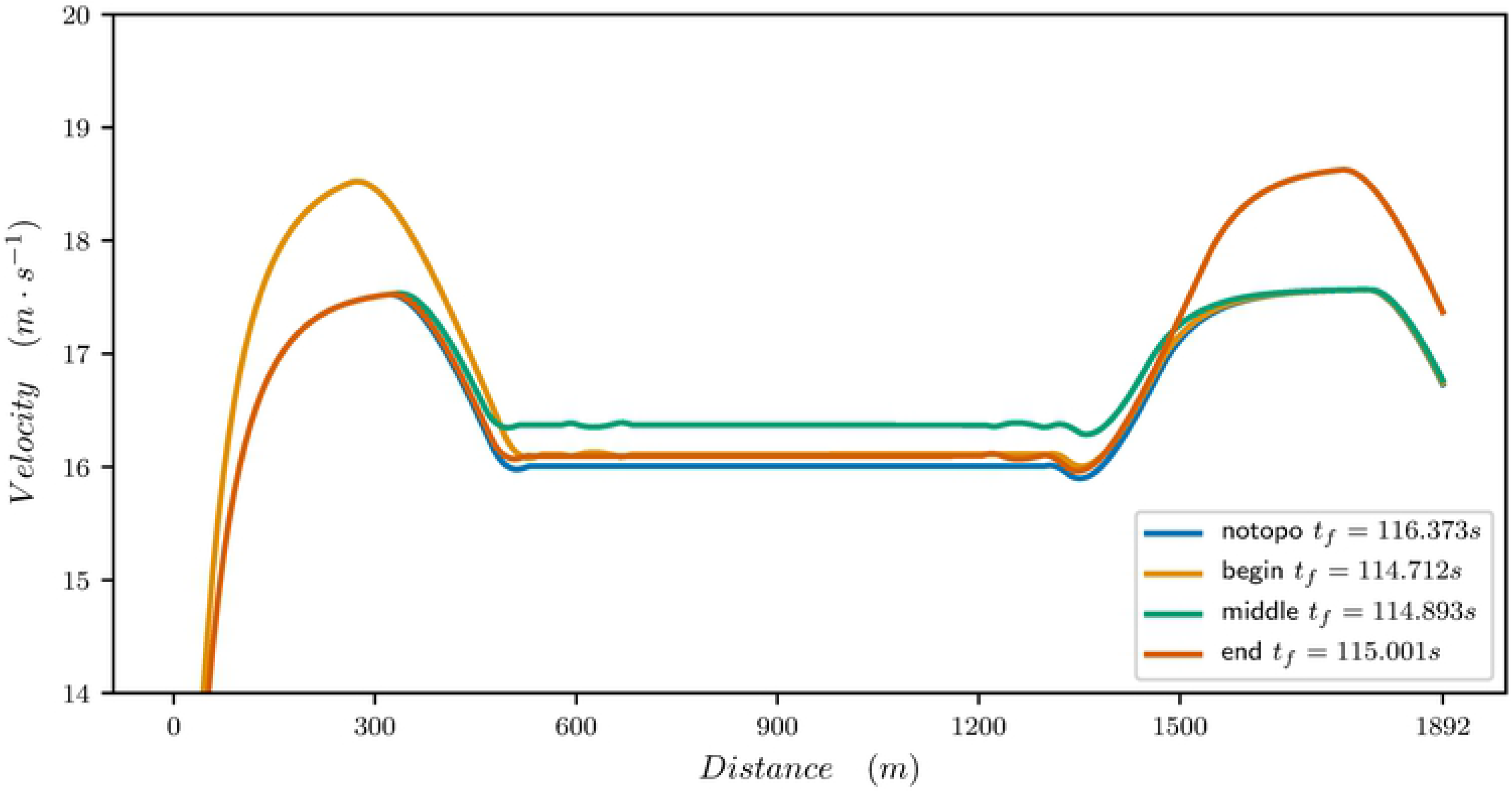
Optimal velocity profile for a 1900m: flat straight track (blue), +3% ramp down for 630m at the beginning of the race (orange), +3% ramp for 630m at the middle of the race (green), +3% slope for 630m at the end of the race (red).

**Fig 12.**
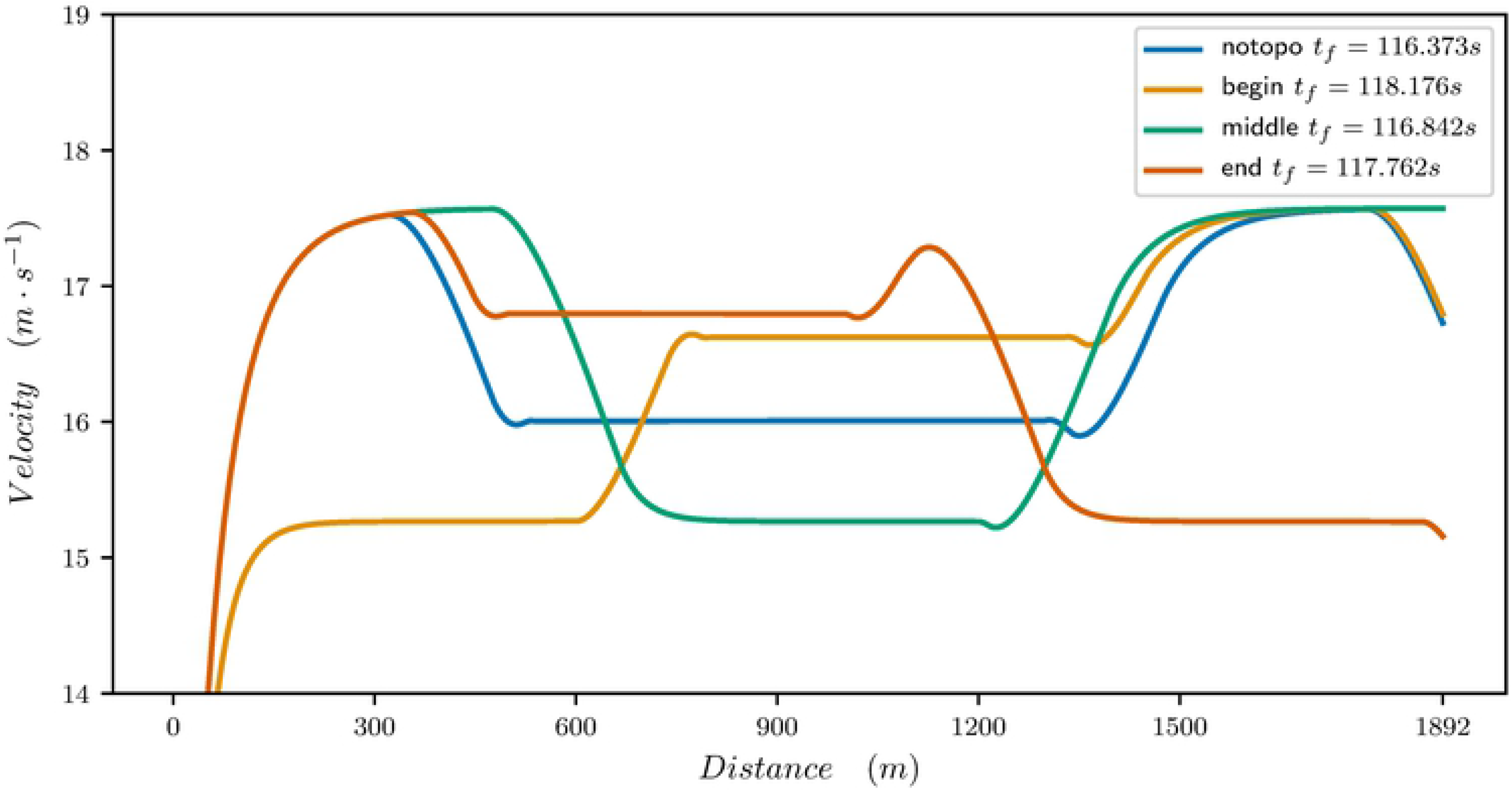
Optimal velocity profile for a 1900m: flat straight track (blue), bend for 630m at the beginning of the race (orange), bend for 630m at the middle of the race (green), bend for 630m at the end of the race (red).

For a slope going up, as illustrated in Fig 10, the best time is obtained when the slope is at the end of the race. Indeed, a good horse can still provide a strong effort at the end of the race, even if he is tired. If the slope is at the beginning, it has a strong effect on the velocity which cannot reach its maximum value. In the middle of the race, the slope reduces the mean velocity and therefore the final time.

For a ramp going down, as illustrated in Fig 11, it is the opposite: the best time is obtained when the ramp is at the beginning of the race. Indeed the horse speeds up more easily and more quickly.

For the bend, as illustrated in Fig 12, the best time is obtained when the bend is at the middle of the race. This is where it has the smallest decrease on the velocity profile. At the beginning, it prevents the horse from reaching its maximal speed. Of course, here the effect is exaggerated with respect to a real race because the bend is long but it yields the general flavour. Similarly, at the end, it prevents the horse from sprinting.

Therefore, we have seen that the pacing strategy has to be analyzed with respect to the changes of slopes, ramps or bends in order to optimize the horse effort. For a given track, it is very likely that the turnpike theory of [19] should yield more precise and detailed analytical estimates of the increase of decrease of velocity.

## Conclusion

Thanks to precise velocity data obtained on different races, we are able to set a mathematical model which provides the optimal pacing strategy for horses on a fixed distance. It relies on both mechanical, energetic considerations and motor control. The process consists in identifying the physiological parameters of the horse from the data. Then the optimal control problem provides information on the pacing. We see that horses have to start strongly and reach a maximal velocity. The velocity decreases in the bends; when going out of the bend, the horse can speed up again and our model can quantify exactly how and when. The horse that slows down the least at the end of the race is the one that wins the race. We understand from the optimal control problem that this slow down is related to the anaerobic supply, the *V̇ O*2_*max*_ and the ability to maintain maximal force at the end of the race. Therefore, horses that have a tendency to slow down too much at the end of the race should put less force at the beginning and slow down slightly through the whole race in order to have the ability to maintain velocity at the end.

From our simulations, we are also able to get information on the *V̇ O*2 profile, such as when steady state *V̇ O*2 is reached, when the decay of *V̇ O*2 starts. The ability to maintain a high *V̇ O*2 for a long time is related to the ability to maintain velocity. The *V̇ O*2 starts to decrease when the residual anaerobic energy is too low, and this corresponds to the optimal time to launch the sprint for a long race.

We also understand better the effects of altitude and the bends and find that they are not local effects producing a local perturbation in the pacing strategy, but on the contrary have a global effect on the whole race. Therefore, a good knowledge of the track and training are crucial to adapt the global pacing strategy rather than slowing down because of bends or slopes.

Future works will be devoted to taking additionally into account drafting and the horse psychology since an alternative strategy can be to stay behind to save energy and overtake in the last straight [25].

## Acknowledgments

The authors acknowledge support from the LabEx AMIES (ANR-10-LABX-0002-01) of Université Grenoble Alpes, within the program “Investissements d’Avenir” (ANR-15-IDEX-0002) operated by the French National Research Agency (ANR).

The authors also wish to thank France Galop and Mc Lloyd for providing the data and for their interest in this work. Finally, they are very grateful to Pierre Martinon for his advice on Bocop.

